# A chemical strategy toward novel brain-penetrant EZH2 inhibitors

**DOI:** 10.1101/2021.06.10.447852

**Authors:** Rui Liang, Daisuke Tomita, Yusuke Sasaki, John Ginn, Mayako Michino, David J. Huggins, Leigh Baxt, Stacia Kargman, Maaz Shahid, Kazuyoshi Aso, Mark Duggan, Andrew W. Stamford, Elisa DeStanchina, Nigel Liverton, Peter T. Meinke, Michael A. Foley, Richard E. Phillips

## Abstract

Aberrant gene-silencing through dysregulation of polycomb protein activity has emerged as an important oncogenic mechanism in cancer, implicating polycomb proteins as important therapeutic targets. Recently, an inhibitor targeting EZH2, the methyltransferase component of PRC2, received FDA approval following promising clinical responses in cancer patients. However, the current array of EZH2 inhibitors have poor brain-penetrance limiting their use in patients with CNS malignancies, a number of which have been shown to be sensitive to EZH2 inhibition. To address this need, we have identified a chemical strategy, based on computational modeling of pyridone-containing EZH2 inhibitor scaffolds, to minimize P-glycoprotein activity and here we report the first brain-penetrant EZH2 inhibitor, TDI-6118 (compound 5). Additionally, in the course of our attempts to optimize this compound we discovered TDI-11904 (compound 21); a novel, highly-potent, and peripherally active EZH2 inhibitor based on a 7 member ring structure.

## Introduction

The characterization of somatic genomes in cancer through efforts such as TCGA (The Cancer Genome Atlas),^1^ has shown chromatin factors are highly mutated in cancer, with up to 40% of cancers bearing at least one mutation in an epigenetic regulator.^2^ This work has generated much interest in targeting epigenetic regulators therapeutically in cancer. One of the most well-characterized epigenetic regulators is EZH2 (enhancer of zeste homologue 2), the methyltransferase (‘writer’) component of the polycomb repressive complex 2 (PRC2).^3^ EZH2 catalyzes the generation of histone-3 lysine 27 methylation (H3K27me), ^4^ an epigenetic mark which is critical for mediating gene repression in many contexts including development. ^5^ A wealth of studies have implicated an oncogenic function for EZH2 through its ability to repress tumor suppressor genes^6^ stimulating many EZH2 drug discovery programs.^7,8,9^ For example, certain cancers such as lymphoma harbor recurrent gain-of-function mutations in EZH2, leading to aberrant gene repression through enhanced production of H3K27 trimethylation (H3K27me3).^10,11^ Additionally, tumors which bear mutations in the SWI-SNF (SWItch/Sucrose Non-Fermentable) complex including sarcomas are frequently sensitive to EZH2 inhibition,^12,13,14^ due to an evolutionary conserved synthetic lethal relationship between SWI-SNF and polycomb.^15,16^

Validation of EZH2 as a bona fide therapeutic target in cancer was recently achieved with encouraging clinical responses to tazemetostat in patient trials leading to FDA approval in epithelioid sarcoma and follicular lymphoma.^17,18^ Although this and other drug discovery programs have been effective in generating EZH2 inhibitors for human clinical trials, there remains a need for novel chemical matter to address other specific contexts in oncology and beyond. One important area of unmet need is CNS malignancies. ^19^ Twenty percent of all patients with cancer will develop brain metastases^20,21^ and several of the cancer-types which have a propensity to metastasize to the brain have been shown to be effectively treated with EZH2 inhibition in pre-clinical cancer models, including small cell lung cancer, ^22^ specific breast adenocarcinoma subtypes^23^ and melanoma.^24,25^ Furthermore, malignant brain tumors are the leading cause of cancer-related mortality in children.^26^ Notably, several pediatric brain tumors are characterized by a reprogrammed epigenomic landscape in which EZH2 is essential for tumor maintenance including midline glioma, ^27, 28^ ependymoma ^29^ and atypical rhabdoid tumors of the brain^30, 31^ all of which lack effective therapies. Thus, a brain-penetrant EZH2 inhibitor could be of great utility in oncology.

Many successful case studies have been reported in the literature where brain penetration is enhanced by minimizing P-glycoprotein (P-gp) efflux transport and increasing passive permeability. ^32^ Hydrogen bond donor count is one key physicochemical property which influences P-gp substrate activity,^33,34^ and reducing the number of hydrogen bond donors has been shown as an effective strategy to lower efflux transport.^35,36,37,38^ While EZH2 inhibitors reported in the literature possess an overall broad diversity of structures, most share a common, conserved pyridone which is critical for engaging and inhibiting EZH2 enzymatic activity [Tazemetostat (**1**),^7^ PF-06726304 (**2**),^39^ PF-06821497 (**3**),^40^ CPI-1205 (**4**),^41^ (Fig. 1)]. It has been shown that pyridone containing EZH2 inhibitors are substrates of efflux transporters which severely restrict their distribution into the brain.^42^ We hypothesized that the common pyridone motif represents a good substrate for efflux transporters due to the presence of a hydrogen bond donor NH group. We envisioned that masking the hydrogen bond donor NH group in the pyridone could produce compounds that maintain a favorable potency profile and possess a reduced efflux ratio leading to small molecule inhibitors with improved brain penetration properties.

**Figure 1.**
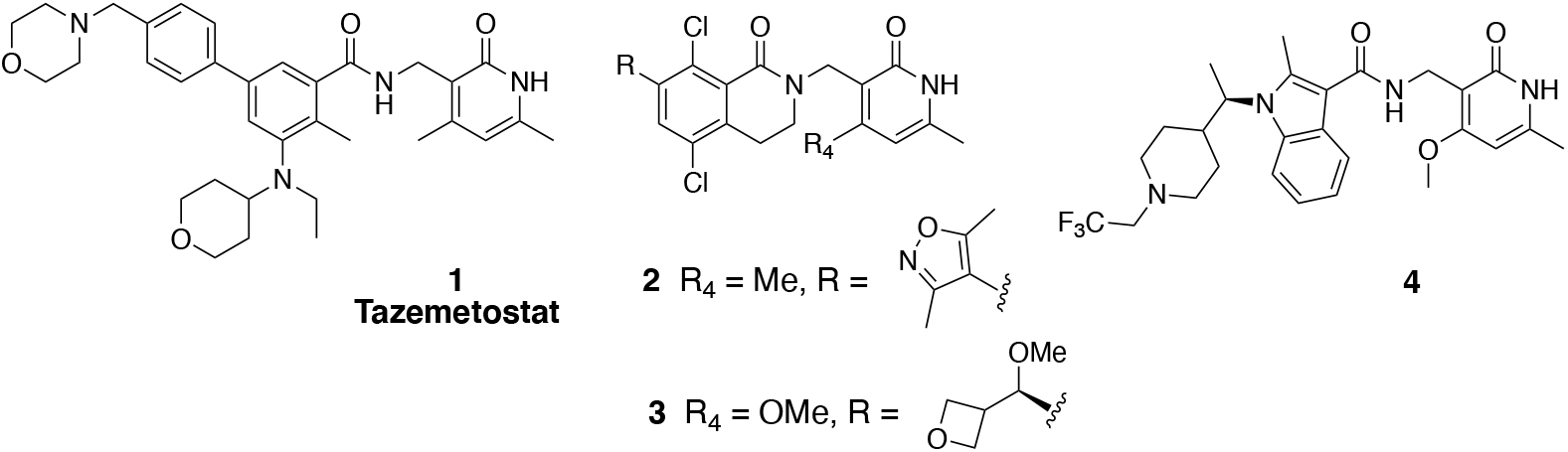
EZH2 inhibitors reported in the literature.

## Results and Discussion

We initiated our medicinal chemistry campaign with an analysis of the crystal structure of Polycomb repressive complex 2 (PRC2) bound to a pyridone-containing inhibitor (PDB:5IJ7). ^43^ The pyridone moiety forms two hydrogen-bond interactions with the backbone of Trp624 from the conserved GXG motif of the SET domain and a water-mediated interaction with the sidechain of Asn688. Calculation of hydration site thermodynamics using WaterMap^44^ suggested that the bridging water is only weakly bound (ΔG=3.09 kcal/mol; ΔH=-1.36; -TΔS=4.45), indicating that it could be displaced without a large desolvation penalty (Fig. 2a). Methylation of the pyridone N-H would remove one of the two hydrogen-bond interactions with Trp624 and displace the bridging water, as predicted by the docking model of a methylated analog (Fig. 2b). Our analysis and docking model suggested that methylation of the free NH in the pyridone would be tolerated and that binding energy could be recovered through modification on the other parts of the molecule such as the 4-position of the pyridone, as shown by others.^45^

**Figure 2.**
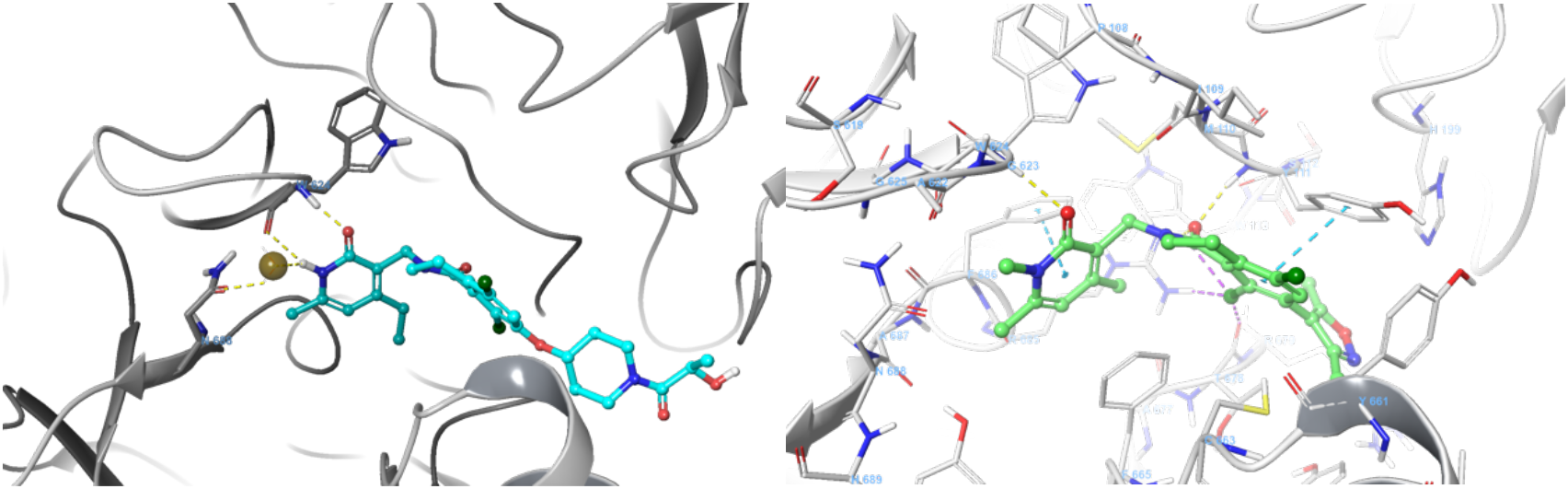
a) Watermap calculation showed de-stabilizing energy at the bridging water hydration site. b) Docking model of methylated pyridone compound **5** in EZH2 (PDB: 5IJ7)

Encouraged by this analysis, we selected a set of disclosed pyridone series EZH2 inhibitors and methylated the NH group in the conserved dimethylpyridone substituent, reducing the number of hydrogen bond donors and aiming to increase brain penetration. While methylation on the pyridone resulted in an expected reduction in biochemical potency we were still able to identify compounds with low nM activity against EZH2. More importantly, several of these modified EZH2 inhibitors exhibited improved MDR efflux ratios compared to the parent N-H pyridone, supporting our strategy of N-methylation on the pyridone. One of these compounds, **5**, exhibited an IC_50_ of 14 nM in our biochemical assay, a modest drop in potency relative to the original compound **2**. Compound **5** retained moderate potency in the cellular EZH2 inhibition assay (flow cytometric quantitation of H3K27me3) (Table 1) while displaying a very favorable MDR ratio of 0.95 and good permeability, representing a significant improvement relative to compound **2**. Furthermore, the concentration ratio of unbound drug in brain to blood (Kp,uu = 0.29) obtained from *in vivo* study in rats demonstrated that compound **5** is brain penetrant *in vivo* and provided proof-of principle that N-methylation of the pyridone group was sufficient to confer brain penetrance. Unfortunately, the instability toward both human and mouse liver microsomes, as well as, modest *in vitro* anti-proliferation effect compared to parent compound **2** limited the utility of **5** in follow up studies. To utilize this novel class of brain penetrant EZH2 inhibitor for *in vivo* studies, we sought to further improve the compound’s cell potency and liver microsomal stability.

**Table 1.** Methylation of dimethylpyridone to improve BBB

|  | **2** | **5** |
| --- | --- | --- |
| EZH2 IC<sub>50</sub> (nM) | 1.0 | 14 |
| Cellular IC<sub>50</sub> (H3K27me3) (nM) | 16 | 580 |
| Karpas-422 proliferation (nM) | 122 | 4680 |
| HLM/MLM (μL/min/mg) | 210/253 | 449/>768 |
| Cl<sub>total</sub> (μL/min/g) | 35 | 100 |
| AUC<sub>po</sub> (h.ng/mL)/Bioavailability | 67.1/13.7% | 5.8/3.5% |
| MDCK/MDR1 permeability(nm/s)<br>AB, efflux ratio | 37, 4.8 | 105, 0.95 |
| K<sub>p,uu</sub> | < 0.003 | 0.29 |

Our approach to improve potency and metabolic stability was initially guided by literature reports exploring substitution at the 4 position of the pyridone.^45^ These showed that the 4-position of the pyridone ring can accommodate a larger hydrophobic group, which is involved with Van der Waals contacts with hydrophobic residues on the protein. Changes at this position can have a pronounced effect on EZH2 enzyme activity ^45^ and we reasoned that the potency loss resulting from N-methylation could be recovered by introducing favorable substituents at the 4-position of the pyridone in our brain penetrant molecules. We then prepared several analogs related to **5** in an effort to improve EZH2 inhibition activity (Table 2). When we incorporated increasingly more hydrophobic substituents (compounds **6a-c**), we found that the MDR ratios were maintained at a desirable level. However, their biochemical potency dropped dramatically. We did observe moderate improvement in human microsome stability; however, stability against mouse liver microsome remained very low. A methoxy group was incorporated to reduced lipophilicity, anticipating improved microsomal stability predicted by lower AlogP value. Compound **6d** displayed a moderate human liver microsome (HLM) increase, however mouse liver microsomes (MLM), biochemical and cellular potency were similar to compound **5**. Surprisingly, compound **6d** exhibited a greatly increased MDR ratio suggesting a very narrow SAR space to maintain desirable permeability characteristics. A similar observation was made by incorporation of a 4-thiomethyl pyridone, recently reported to enhance EZH2 potency of compound **4** (Fig. 1) with increased residence time,^46^ to provide compound **6e** which exhibited biochemical and cellular potency similar to parent compound **2**. Therefore, while the 4-SMe group was able to restore the potency lost due to N-methylation in compound **5**, the desirable brain-penetrating properties gained through N-methylation were lost.

**Table 2.** SAR on 4-position of the pyridone

| Compound | R4 | EZH2<br>IC<sub>50</sub><br>(nM) | Karpas-422<br>Proliferation<br>(nM) | HLM/MLM<br>(μL/min/mg) | AlogP | MDCK/MDR1<br>permeability(nm/s)<br>AB, BA/AB |
| --- | --- | --- | --- | --- | --- | --- |
| **5** | Me | 14 | 4680 | 449/>768 | 4.4 | 105, 0.95 |
| **6a** | Et | 82 | - | >768/>768 | 4.9 | 110, 0.72 |
| **6b** | *c*-Pr | 302 | - | 490/>768 | 4.7 | 52, 1.5 |
| **6c** | CF<sub>3</sub> | >1000 | - | 287/592 | 4.9 | 49, 2.4 |
| **6d** | OMe | 11.6 | 2950 | 117/656 | 3.9 | 10, 23 |
| **6e** | SMe | 1.3 | 632 | 235/796 | 4.5 | 3, 110 |

Further effort to improve the potency and metabolic stability of N-methyl pyridone-containing EZH2 inhibitors led us to initiate a medicinal chemistry effort to identify a new class of scaffold. Kung et al. showed that a seven-membered-ring analog related to compound **2** exhibited comparable potency and metabolic stability.^39^ Our modeling studies based on crystal data (Fig. 3) showed that there is overall good overlay between 6- and 7-membered-ring structures. The expanded 7-membered ring occupied a similar space with potential for additional interactions with the protein. Guided by our computational docking model, we then prepared several analogs related to compound **2** (Table 3) with progressively larger substituents R1 at the aniline nitrogen. We chose to use the 4-OMe pyridone because it imparted higher microsomal stability (likely due to lower lipophilicity), as well as slightly more potent EZH2 biochemical activity. Our first set of compounds included **7** (R_1_ = H) and **8** (R_1_ = Me), which both exhibited subnanomolar potency in the biochemical assay and were as potent as the parent 6-membered ring compound **2**. The enhanced biochemical potency translated into similar or better potency in cellular H3K27me3 assays and antiproliferation assays (Table 3). Compounds **7** and **8** also displayed improvements in HLM/MLM stability parameters relative to the analogous 6-membered-ring compound **2**, presumably attributed to the decrease in AlogP values by incorporating the additional nitrogen in the 7-membered ring. Substitution of the aniline nitrogen with a larger ethyl group, compound **9**, displayed dramatic attenuation of biochemical potency. It is unclear whether the potency drop was caused by the conformation change in the 7-membered ring due to steric repulsion between the ethyl group and the Cl group, or simply steric hinderance in the binding pocket to accommodate the ethyl group that is not predicted by our docking model. By removing the Cl group, we anticipated that a less restricted conformation would be populated due to the relief of steric hinderance, and at the same time, further reduction of AlogP will bring the HLM/MLM stability to a favorable level. We then prepared analogous des-Cl compounds **10** (R_1_ = H) and **11** (R_1_ = Me). Compound **10** displayed a modest loss in biochemical potency and antiproliferation efficacy, which is consistent with the effect of Cl substituent in the 6-membered ring reported in the literature.^39^ Interestingly, the decrease in AlogP resulted in significant improvement in both HLM and MLM stability. Unexpectedly, the ethyl substituted compound **11** exhibited more significant potency loss and we decided not to pursue more extensive SAR at this position.

**Table 3.**
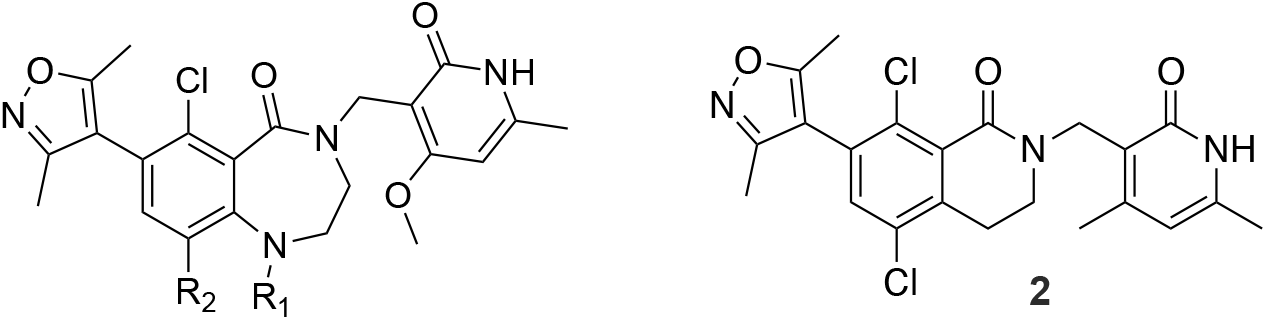
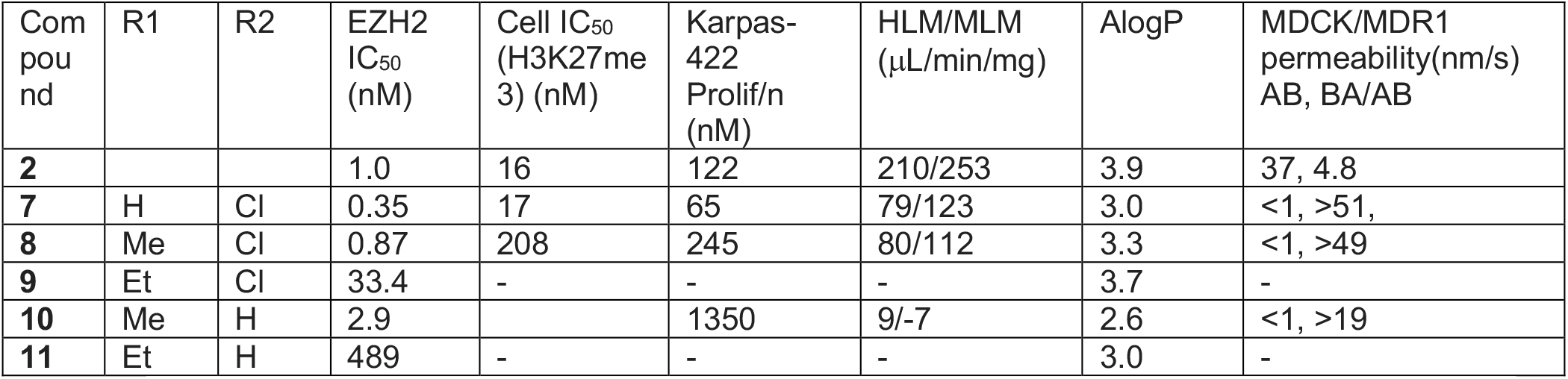
SAR on 7-membered-ring scaffold

| Compound | R1 | R2 | EZH2 IC<sub>50</sub> (nM) | Cell IC<sub>50</sub> (H3K27me3) (nM) | Karpas-422 Prolif/n (nM) | HLM/MLM (μL/min/mg) | AlogP | MDCK/MDR1 permeability(nm/s) AB, BA/AB |
| --- | --- | --- | --- | --- | --- | --- | --- | --- |
| **2** |  |  | 1.0 | 16 | 122 | 210/253 | 3.9 | 37, 4.8 |
| **7** | H | Cl | 0.35 | 17 | 65 | 79/123 | 3.0 | <1, >51, |
| **8** | Me | Cl | 0.87 | 208 | 245 | 80/112 | 3.3 | <1, >49 |
| **9** | Et | Cl | 33.4 | - | - | - | 3.7 | - |
| **10** | Me | H | 2.9 |  | 1350 | 9/-7 | 2.6 | <1, >19 |
| **11** | Et | H | 489 | - | - | - | 3.0 | - |

**Figure 3.**
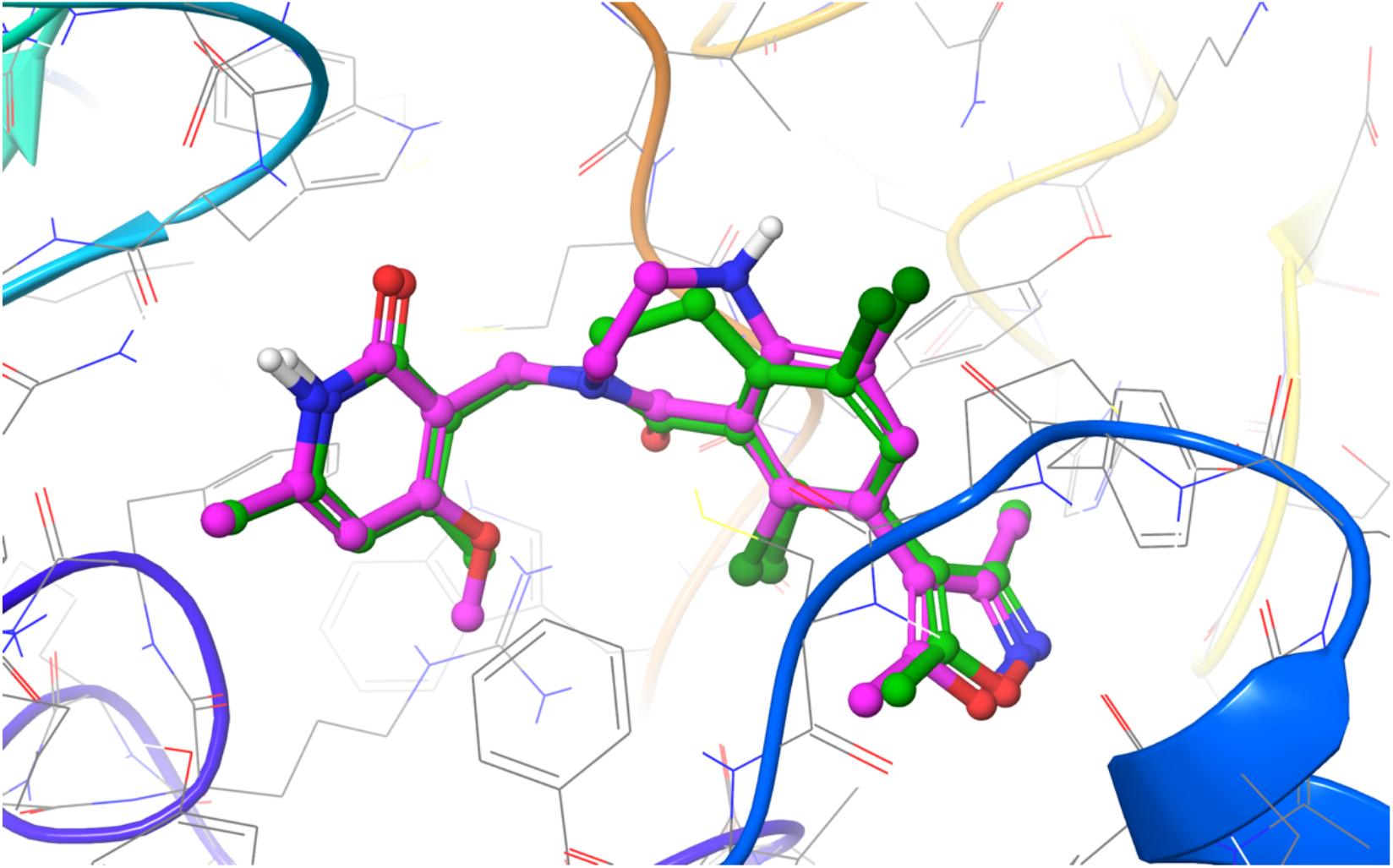
The predicted binding mode between EZH2 and compound **7**, overlaid with the crystallographic binding mode of compound **2** from PDBID 6B3W.^47^ The protein is displayed as cartoon ribbons with atoms in grey wire and the crystallographic binding mode of compound 2 is shown as green balls and sticks. The predicted binding mode of compound **7** is shown as magenta balls and sticks.

The potency and metabolic stability profile of compound **8** led us to explore this substitution pattern in conjunction with SAR at the R_3_ position (Table 4). More importantly, we chose compound **8** over the slightly more potent compound **7** as the starting point because it has one less hydrogen bond donor which would have reduced efflux potential. The substitution explored at the R_3_ position included moieties previously demonstrated by us and others in the 6-membered ring lactam scaffold to display potent biochemical inhibition of EZH2 and/or good metabolic stability.^40^ We were curious to find out if SAR previously established was transferable to the new 7-membered ring lactam scaffold. It was shown that introduction of the sp^3^-carbon atom at this position was tolerated. Substitution of the R_3_ position with an isopropyl group (**12**) in compound **8** resulted in significant loss of potency, suggesting that dimethylisoxazole made similar hydrophobic interactions with the protein as the 6-membered ring lactam scaffold as predicted in the docking model. We then extended the isopropyl group with a branched tetrahydropyran (**13, 15**) to fill the two hydrophobic binding pockets occupied by the dimethylisoxazole, either through a carbon or nitrogen linkage. Both compounds **13** and **15** indeed displayed similar potency to compound **8** but with much lower HLM/MLM stability. Reduction of AlogP through a hydroxyl group (**14**) increased HLM/MLM stability with modest loss of potency. Replacement of dimethylisoxazole with aromatic moieties (**16** - **18**) that we identified in the 6-membered ring lactam series maintained the microsomal stability but displayed significant loss of potency in the new series.

**Table 4:**
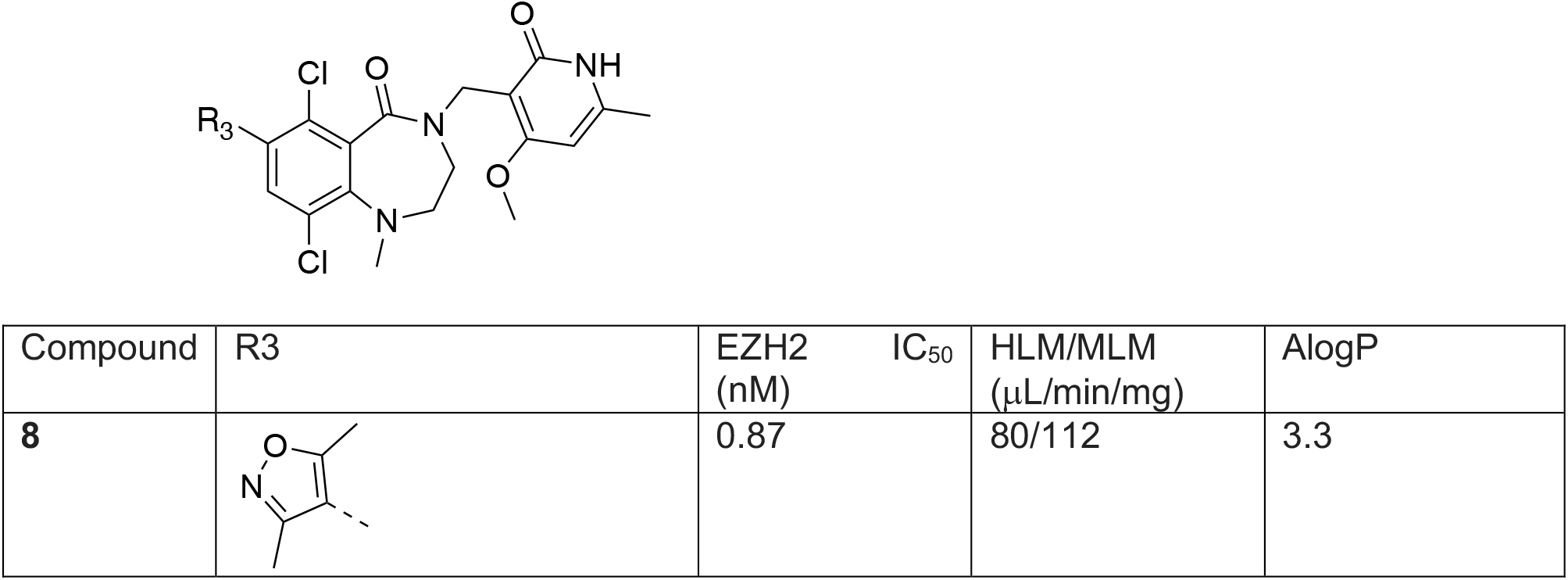
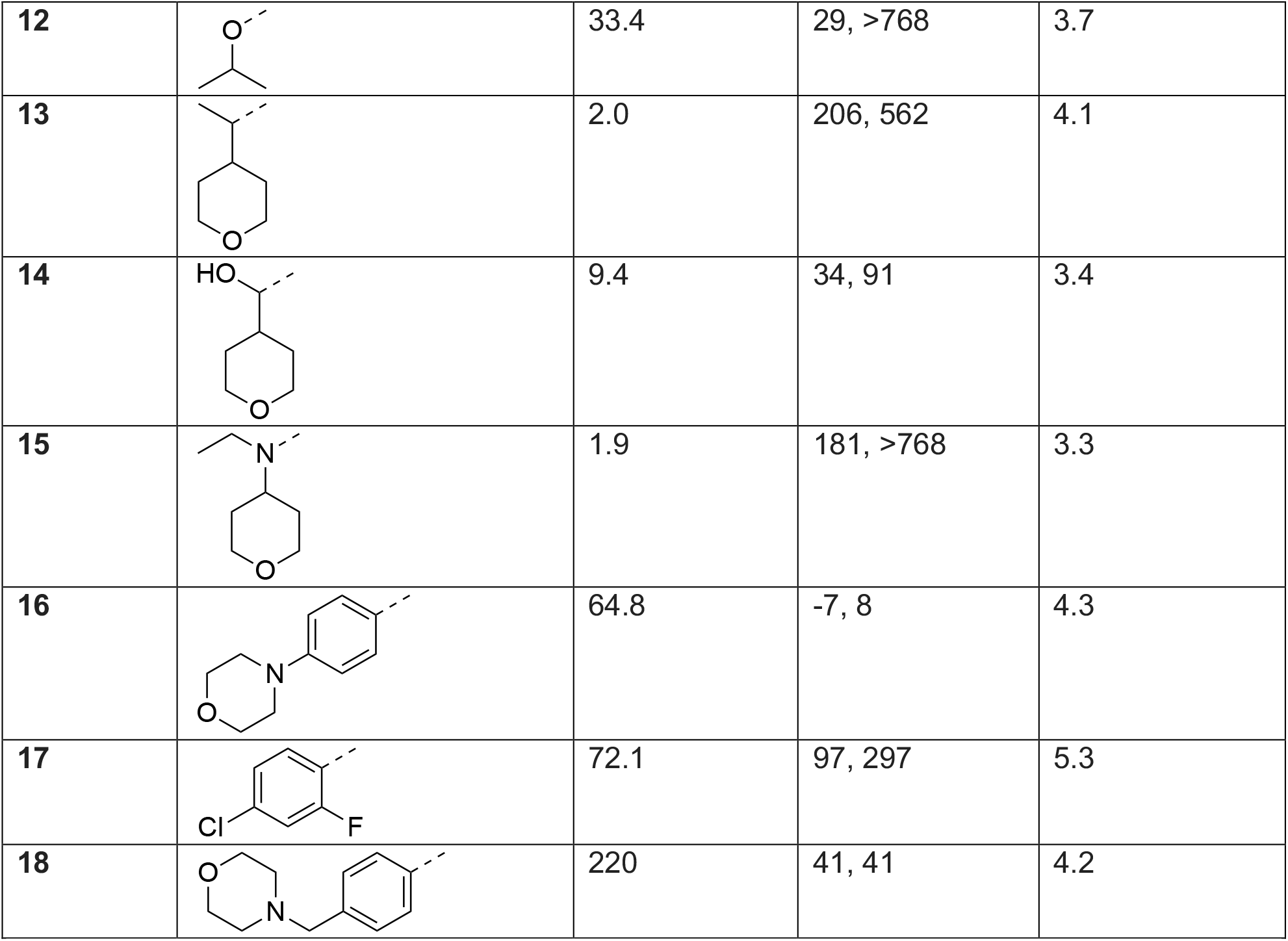
SAR for dimethylisoxazole replacement

| Compound | R3 | EZH2 IC<sub>50</sub> (nM) | HLM/MLM (μL/min/mg) | AlogP |
| --- | --- | --- | --- | --- |
| **8** |  | 0.87 | 80/112 | 3.3 |
| 12 |  | 33.4 | 29, >768 | 3.7 |
| 13 |  | 2.0 | 206, 562 | 4.1 |
| 14 |  | 9.4 | 34, 91 | 3.4 |
| 15 |  | 1.9 | 181, >768 | 3.3 |
| 16 |  | 64.8 | -7, 8 | 4.3 |
| 17 |  | 72.1 | 97, 297 | 5.3 |
| 18 |  | 220 | 41, 41 | 4.2 |

**Table 5.**
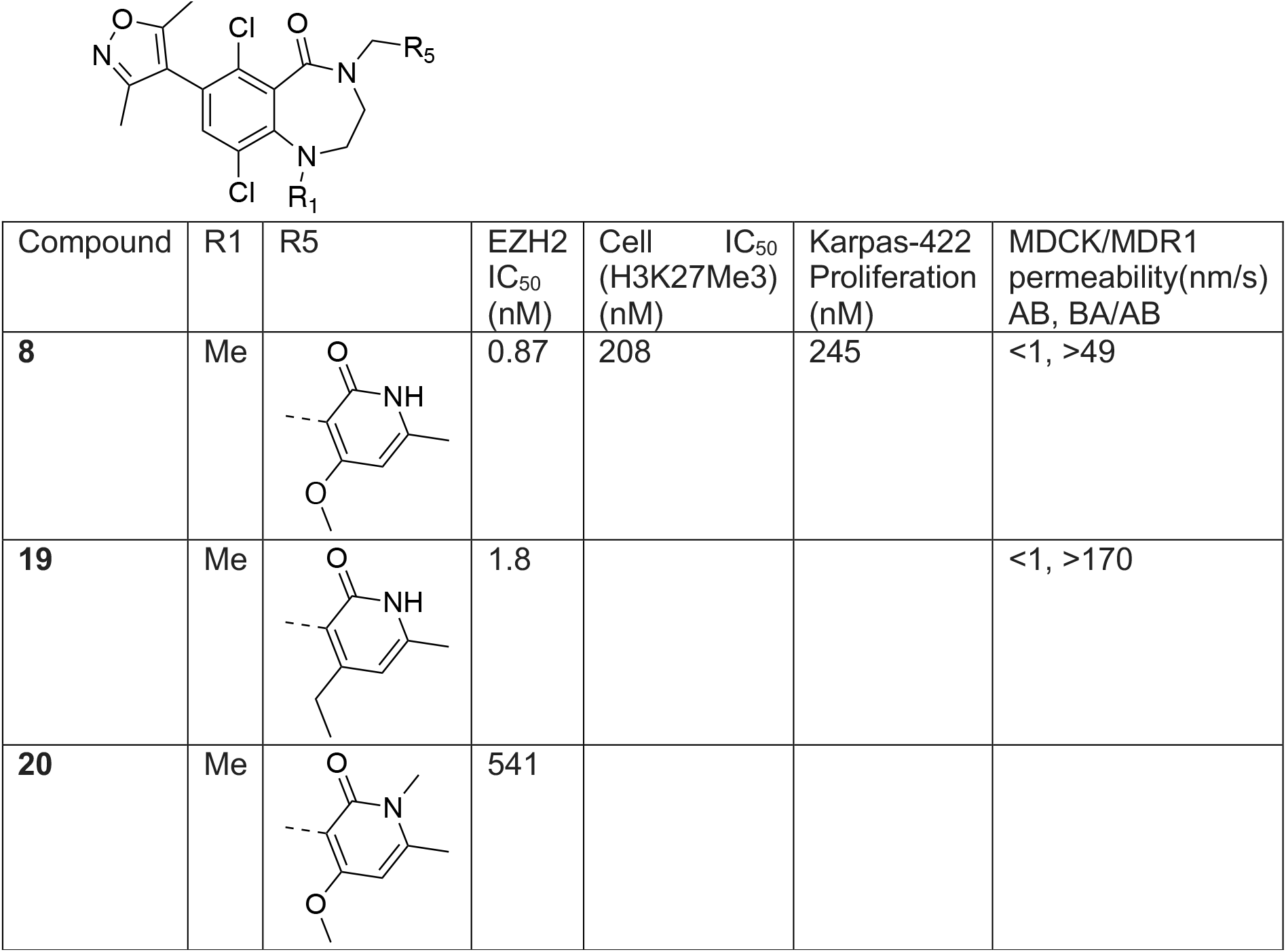
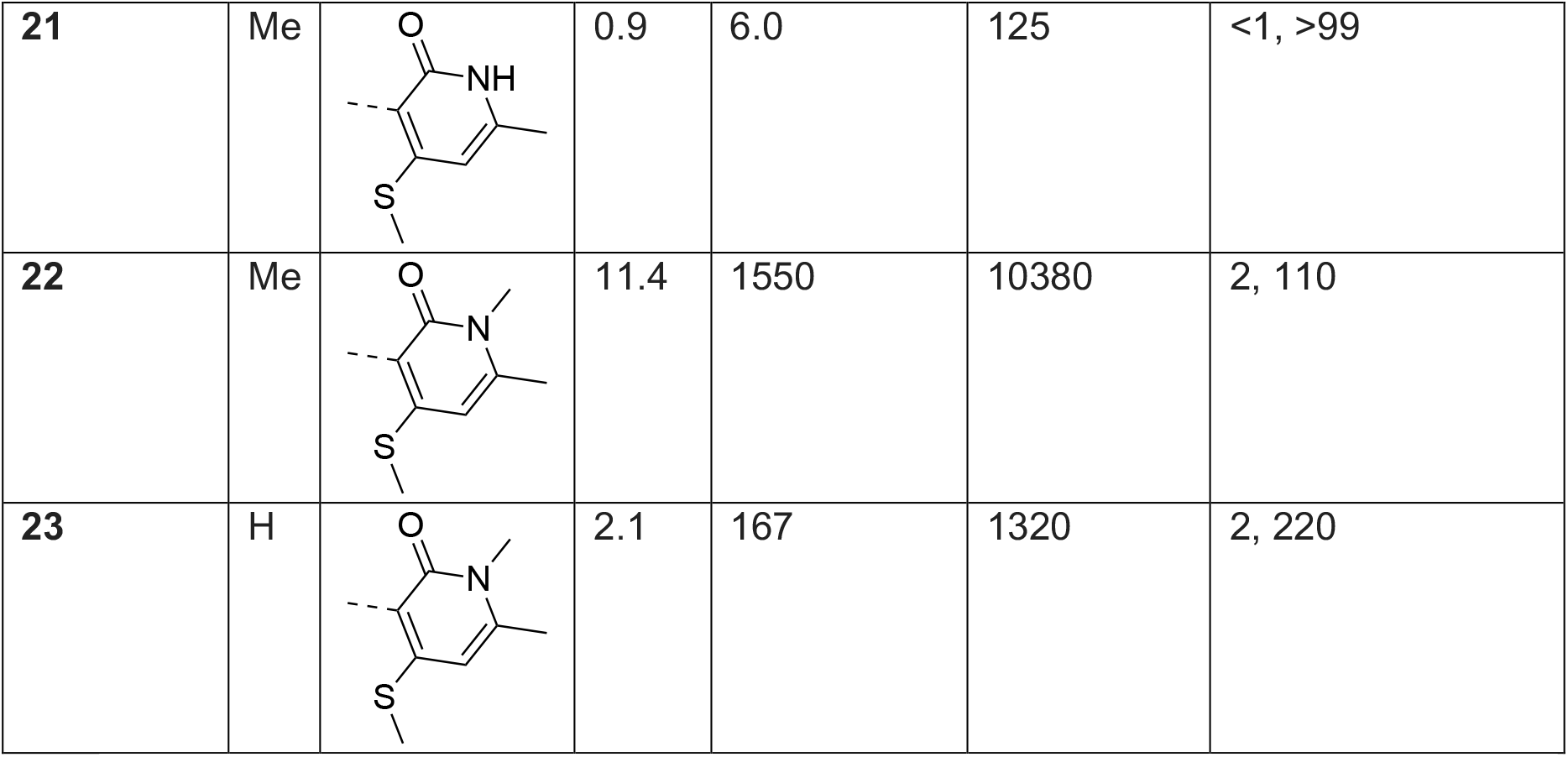
Thiomethyl pyridone analogues with 7-membered ring structure.

| Compound | R1 | R5 | EZH2<br>IC<sub>50</sub><br>(nM) | Cell<br>(H3K27Me3)<br>(nM) | Karpas-422<br>Proliferation<br>(nM) | MDCK/MDR1<br>permeability(nm/s)<br>AB, BA/AB |
| --- | --- | --- | --- | --- | --- | --- |
| **8** | Me |  | 0.87 | 208 | 245 | <1, >49 |
| **19** | Me |  | 1.8 |  |  | <1, >170 |
| **20** | Me |  | 541 |  |  |  |
| **21** | Me |  | 0.9 | 6.0 | 125 | <1, >99 |
| **22** | Me |  | 11.4 | 1550 | 10380 | 2, 110 |
| **23** | H |  | 2.1 | 167 | 1320 | 2, 220 |

To access potent EZH2 inhibitors with favorable brain penetrating properties, we reasoned that the potency profile of the 6,7-scaffold in combination with substitution to either increase lipophilicity or mask hydrogen bond donor functionality would provide a balanced profile to test this drug concept. To this end, we explored replacement of the 4-OMe group with the more lipophilic 4-ethyl substituted pyridone. Unfortunately the resulting compound **19** displayed has no obvious advantage in permeability relative to compound **8** and also appeared to have a similar potency. This was surprising since this substitution has been associated with enhanced EZH2 potency through increasing residence time.^46^ However, the SAR analysis for high affinity compounds is complicated by the enzymatic assay protein concentration of 5 nM resulting in a floor of the assay of 2.5 nM. For example, incorporation of the N-methyl pyridone into compound **20** displayed a greater than 500-fold loss of potency relative to compound **8** but the exact magnitude of the loss is unclear.

During the course of our program, Khanna *et. al* disclosed that incorporating a 4-thiomethyl pyridone in CPI-1205 (**4**) as a key modification significantly increased biochemical and cellular potency due to prolonged residence time, leading to the development of 2^nd^ generation EZH2 inhibitor CPI-0209 with extended target coverage.^49^ We incorporated this finding into our new 7-membered ring structure of EZH2 inhibitor. The resulting compound, compound **21**, exhibited similar biochemical potency relative to compound **8** but superior cellular efficacy (>30-fold improvement), consistent with the literature finding. Importantly, this improvement translated to markedly enhanced potency in anti-proliferation assays in Karpas-422 cells, comparable to those of other reported EZH2 inhibitors including the clinically approved Tazemetostat control. Despite the desirable potency, this compound proved to be a significant MDR substrate and therefore was not suitable for advancement as a brain-penetrant analogue. Nevertheless, the efficacy improvement from the thiomethyl pyridone was very encouraging. Accordingly, we prepared the analogous N-methyl pyridone compound **22**. Although we did observe a significant improvement in biochemical potency for compound **22** versus **20**, this compound still suffered from poor cellular efficacy. Removal of the R1 methyl group (**23**) provided a further improvement in both biochemical and cellular efficacy. Unfortunately, these compounds still displayed high value of MDCK/MDR1 ratio, suggesting that they are good substrates for efflux transporters and would not likely achieve significant brain penetration.

## Conclusion

Utilizing structure-based drug design a novel and highly potent class of 6,7-fused EZH2 inhibitors was identified. This series provided compounds with sub-nanomolar enzymatic activity and potent antiproliferative effects. Enhanced inhibition of cellular trimethylation could be achieved by compound **21** (TDI-11904), derived from incorporation of thiomethyl substitution on the key pyridone moiety (**8** vs **21**). This is in line with reports of improved on target residence time with this substitution on a different scaffold.^46^ Our results demonstrate that this is a general effect that can be applied to different chemical series of EZH2 inhibitors. Additionally, a new strategy to create brain penetrant EZH2 inhibitors through methylation of the conserved pyridone found in many EZH2 inhibitors was described. Compound **5** (TDI-6118) displayed brain penetrating properties both *in vitro* and *in vivo*, and marks the first report of a brain penetrant EZH2 inhibitor. The improved efflux ratio observed with compound **5** could not be maintained with either further modification on the pyridone or with the change to the 7-membered ring lactam structure. However, the enzymatic potency seen by combining the N-methylation with pyridone substitution for increased residence time (*e*.*g*. **22**) suggests that additional structural modifications or alternate cores may provide compounds that balance high-potency (and anti-proliferative effects) with brain penetrance.

### Chemistry

Synthesis of R_1_ methyl-substituted 7-membered lactam was accomplished starting from 2-amino-3,6-dichlorobenzoic acid **24** (Scheme 1). Bromination of compound **24** followed by reductive amination of compound **25** with N-Boc-2-aminoaldehyde **26** afforded bromide carboxylic acid **27**. Treatment of **27** with HCl deprotected the Boc group from the amine moiety, and subsequent cyclization under amide coupling condition generated the lactam intermediate **29**, which upon methylation afforded intermediate **30**.

**Scheme 1.**
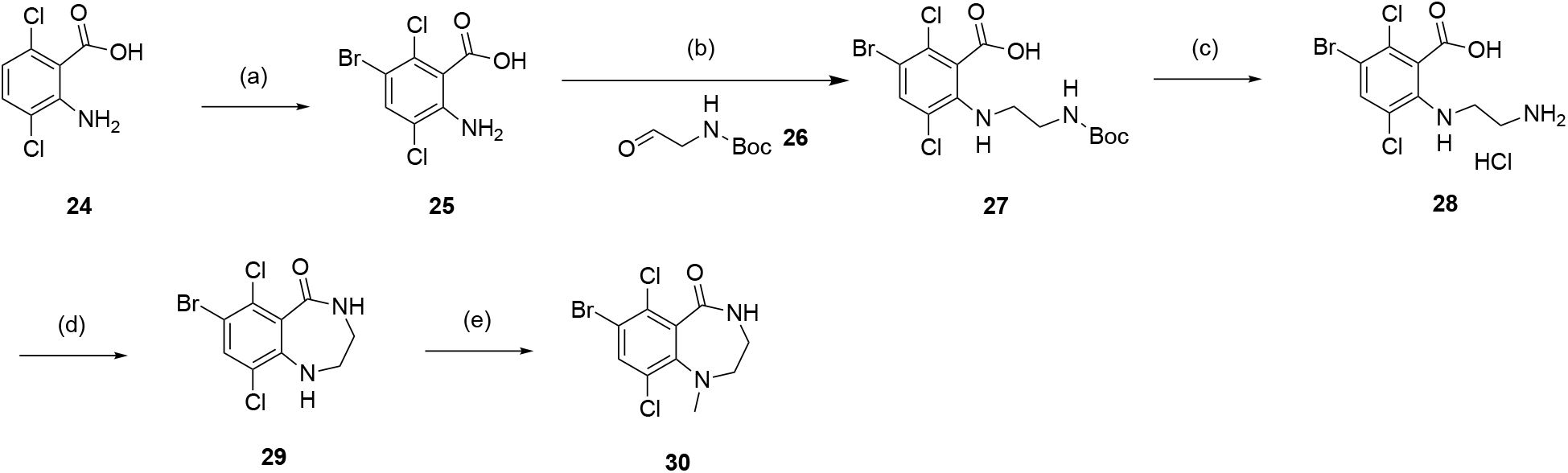
Synthesis of 7-membered ring core bromides **29** and **30**. Reagents and conditions: (a) NBS, DMF, RT; (b) NaBH4, AcOH, DCE, 60°C, 12 h; (c) 4M HCl in dioxane, RT; (d) HOBt, BOP, NMM, DMF, RT, 16 h; (e) TEA, MeI, DMF, 70°C, 16 h.

Benzyloxy-protected chloromethylpyridine intermediate **31** was prepared from commercially available 2-hydroxy-4-methoxy-6-methylpyridine-3-carbonitrile according to literature procedure.^39^ Alkylation of the lactam nitrogen in intermediates **29** and **30** with the chloride intermediate **31** under basic conditions afforded the common intermediates **32** and **33** (Scheme 2). Suzuki coupling with dimethylisoxazole boronic acid **34** followed by the removal of benzyl protecting group with TFA afforded the final compounds **8** and **9**.

**Scheme 2.**
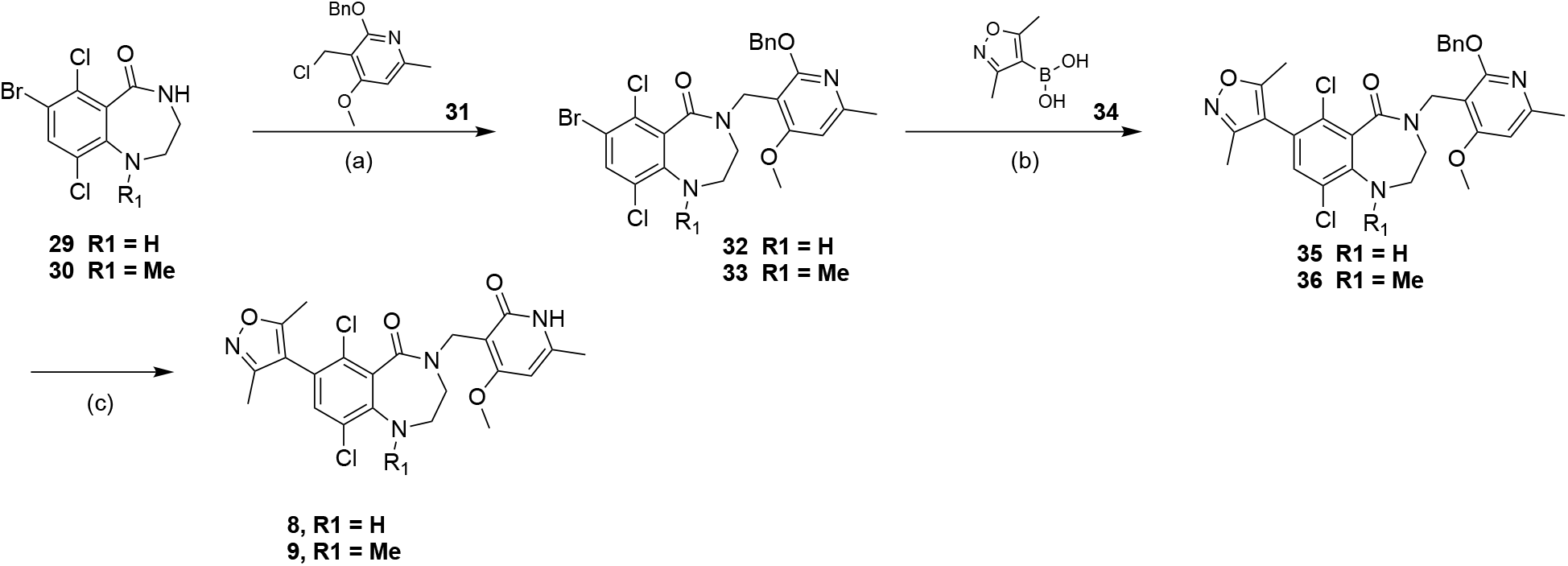
Synthesis of compounds **8** and **9**. Reagents and conditions: (a) *t*-BuOK, DMF for **29**; NaH, DMF for **30**, 40°C, 30 min; (b) Pd(dppf)Cl_2_, K_3_PO_4_, dioxane, H_2_O, 90°C, 2 h; (c) TFA, DCM, RT, 16 h.

Compound **12** was prepared through two sequential alkylations (Scheme 3), one with 2-iodopropane on hydroxyl group of phenol intermediate **38**, the other with chloride intermediate **31** on NH group of amide intermediate **39**. Compounds **13** was accessed through two Grinard reactions (Scheme 4) starting from bromide intermediate **33**. The first Grignard reaction with 4-formyl tetrahydropyran gave secondary alcohol intermediate **41**, which was either subjected to acidic condition to provide compound **14**, or oxidation condition to ketone intermediate **42**. The second Grignard reaction generated tertiary alcohol intermediate **43** which was converted to compound **13** by elimination under acidic conditions.

**Scheme 3.**
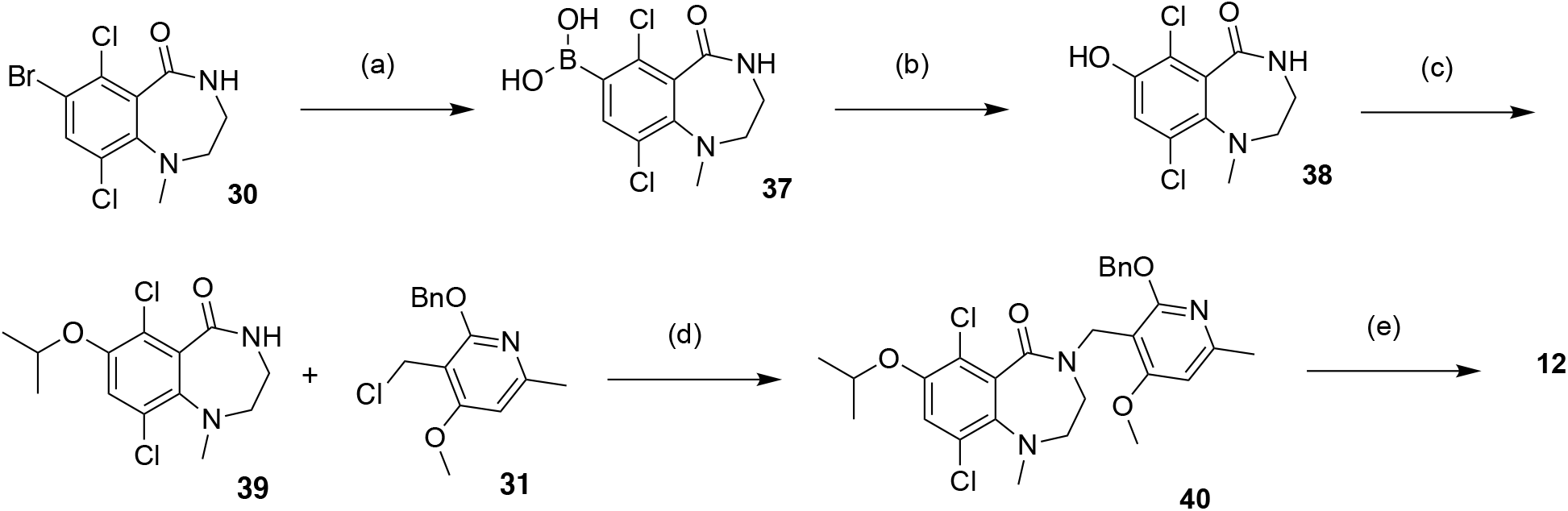
Synthesis of compound **12**. Reagents and conditions: (a) Bis(pinacolato)diborane, Pd(dppf)Cl_2_, KOAc, dioxane, 90°C, 3 h; (b) H_2_O_2_, NaOH, THF, water, 25°C, 2 h; (c) 2-iodopropane, DMF, K_2_CO_3_, 40°C, 2 h; (d) **31**, NaH, DMF, 40°C, 1 h; (e) TFA, DCM, 45°C, 1.5 h.

**Scheme 4.**
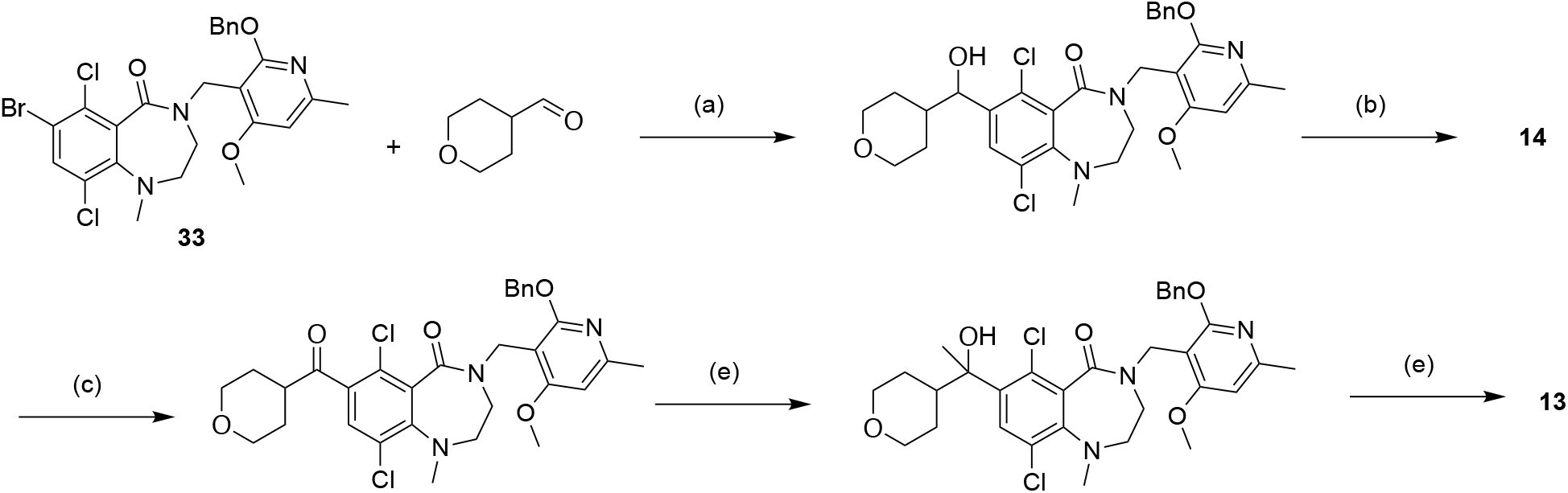
Synthesis of compounds **13-14**. Reagents and conditions: (a) *i*-PrMgCl, THF, −65 to 10°C, 1.5 h; (b) TFA, DCM, 45°C, 1.5 h; (c) Dess-Martin, DCM, 15°C, 12 h; (d) MeMgBr, THF, 0°C, 2 h; (e) Et_3_SiH, TFA, 0 - 45°C, 4 h;

Compound **15** was prepared through reductive amination using amine intermediate **44** (Scheme 5), which was generated by amination of the bromide **33**. Compounds **16-18** with aromatic R3 substitutions (Scheme 6) were synthesized via Suzuki reaction using the corresponding commercial boronic acids **47-49**. Methylation of NH group in pyridone followed the general procedure (Scheme 7).

**Scheme 5.**
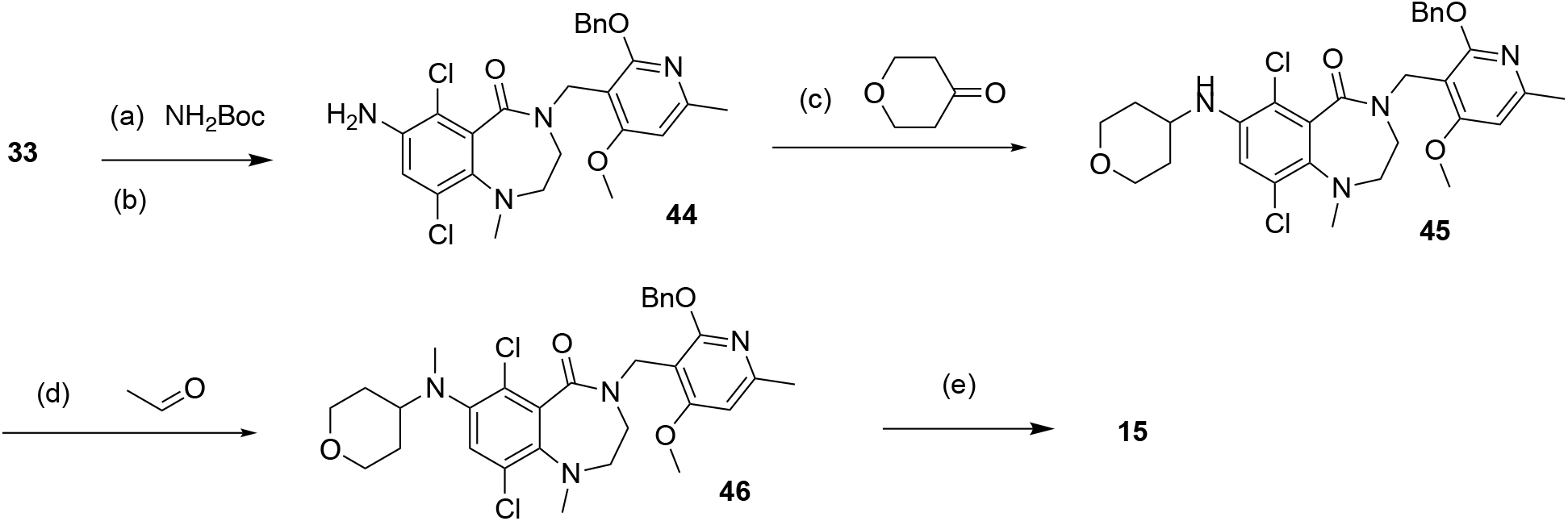
Synthesis of compounds **15**. Reagents and conditions: (a) Pd(dba)_2_, xantphos, Cs_2_CO_3_, dioxane, 110°C, 2 h; (b) HCl/EtOAC, 15°C, 2 h; (c) Tetrahydropyran-4-one, tetramethylammonium, triacetoxyboranuide, TFA, DCM, 15°C, 2 h; (d) Acetaldehyde, tetramethylammonium, triacetoxyboranuide, TFA, DCM, 15°C, 2 h; (e) TFA, DCM, 45°C, 2 h.

**Scheme 6.**
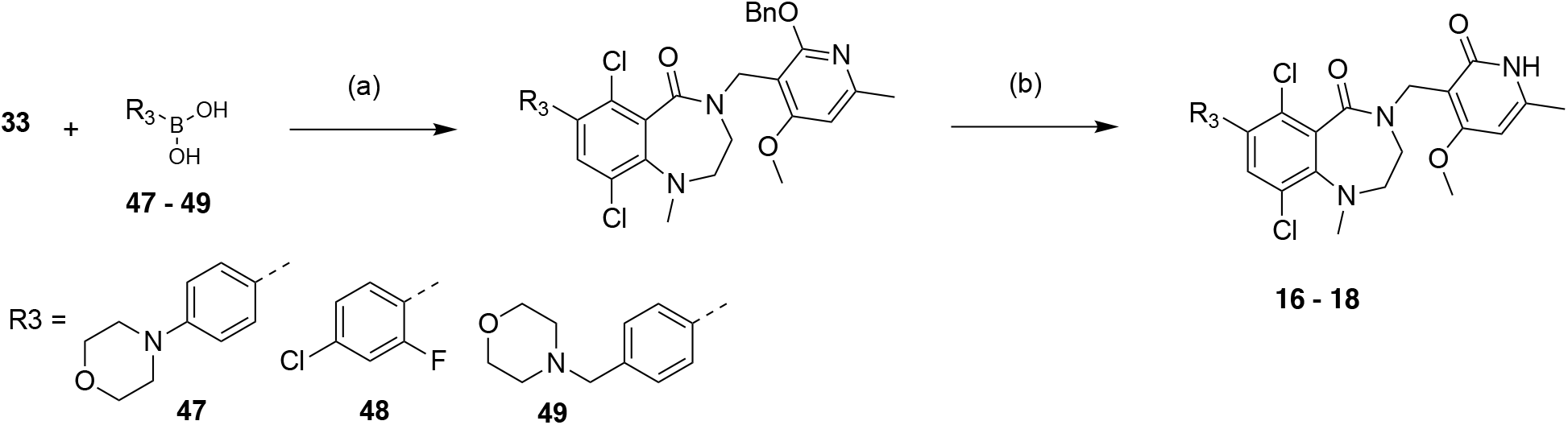
Synthesis of compounds **16-18**. Reagents and conditions: (a) Pd(dppf)Cl_2_, K_3_PO_4_ or CsF, dioxane, H_2_O, 90°C, 2 h; (b) TFA, DCM, RT, 16 h.

**Scheme 7.**
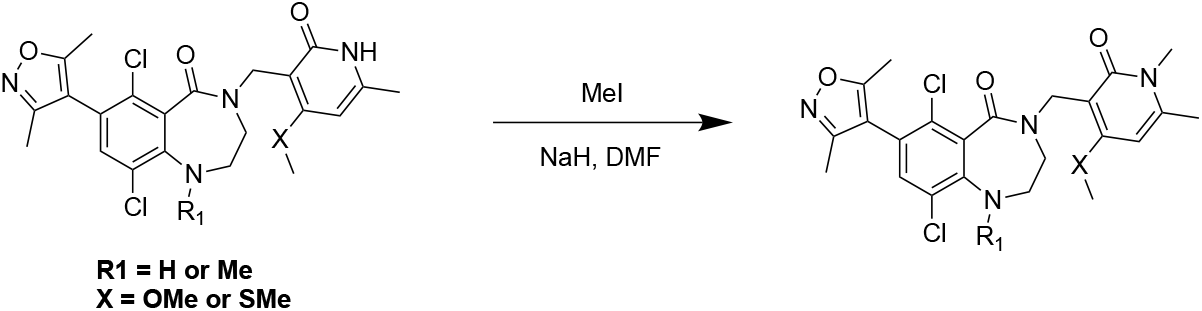
General procedure for N-methylation of pyridone. Reagents and conditions: NaH, MeI, DMF, 0-20°C for 12 h.

## Acknowledgements

The authors gratefully acknowledge the support to the project generously provided by the Tri-Institutional Therapeutics Discovery Institute (TDI), a 501(c)(3) organization. TDI receives financial support from Takeda Pharmaceutical Company, TDI’s parent institutes (Memorial Sloan Kettering Cancer Center, The Rockefeller University and Weill Cornell Medicine) and from a generous contribution from Mr. Lewis Sanders and other philanthropic sources. RP is supported by the MSK IDEA Award and the Center for Experimental Therapeutics, The Cure Starts Now and Team Jack Foundation.

