## Supplemental Information for "A chemical strategy toward novel brain-penetrant EZH2 inhibitors"

#### Contents

General Information

Chemistry

Biology

- *In Vitro* Enzymatic Assay
- Cell Growth Inhibition Assay
- H3K27me3 Flow Cytometry

#### General Information

$^{1}$H NMR spectra and $^{13}$C NMR spectra were recorded on a Bruker 400 MHz spectrometer. Spectra are referenced to residual chloroform (d 7.26, $^{1}$H), DMSO (d 2.54, $^{1}$H) or methanol (d 3.34, $^{1}$H) unless otherwise noted. Chemical shifts are reported in ppm (d); multiplicities are indicated by s (singlet), d (doublet), t (triplet), q (quartet), quint (quintet), sext (sextet), m (multiplet) and br (broad). Coupling constants, *J*, are reported in Hertz. Silica gel chromatography was performed using a Teledyne Isco CombiFlash® Rf+ instrument using Hi-Purit Silica Flash Cartridges (National Chromatography Inco) or RediSep Rf Gold C18 Cartridges (Teledyne Isco). Analytical HPLC was performed on a Waters ACQUITY UPLC with a photodiode array detector using and a Waters ACQUITY BEH Shield RPC18 (2.1 × 50 mm, 1.7 µm) column. Analytical LCMS was performed on a Waters ACQUITY UPLC with a Waters 3100 mass detector. Chiral HPLC was performed on a Waters Alliance e2695 with a photodiode array detector using Daicel Chiralpak® AD-H, Chiralpak® IA, Chiralpak® IB, Chiralpak® IC, Chiralcel® OD-H or Chiralcel® OJ-H columns. Optical rotations were obtained on a Jasco P-2000 digital polarimeter and are reported as [α]D T temperature (T), concentration (c = g/100 mL) and solvent. Commercially available reagents and solvents were used as received unless otherwise indicated.

### Chemistry

**Synthesis of 6,9-dichloro-7-(3,5-dimethylisoxazol-4-yl)-4-((4-methoxy-6-methyl-2-oxo-1,2-dihydropyridin-3-yl)methyl)-1,2,3,4-tetrahydro-5*H*-benzo[*e*][1,4]diazepin-5-one (8) and 6,9-dichloro-7-(3,5-dimethylisoxazol-4-yl)-4-((4-methoxy-6-methyl-2-oxo-1,2-dihydropyridin-3-yl)methyl)-1-methyl-1,2,3,4-tetrahydro-5*H*-benzo[*e*][1,4]diazepin-5-one (9):**

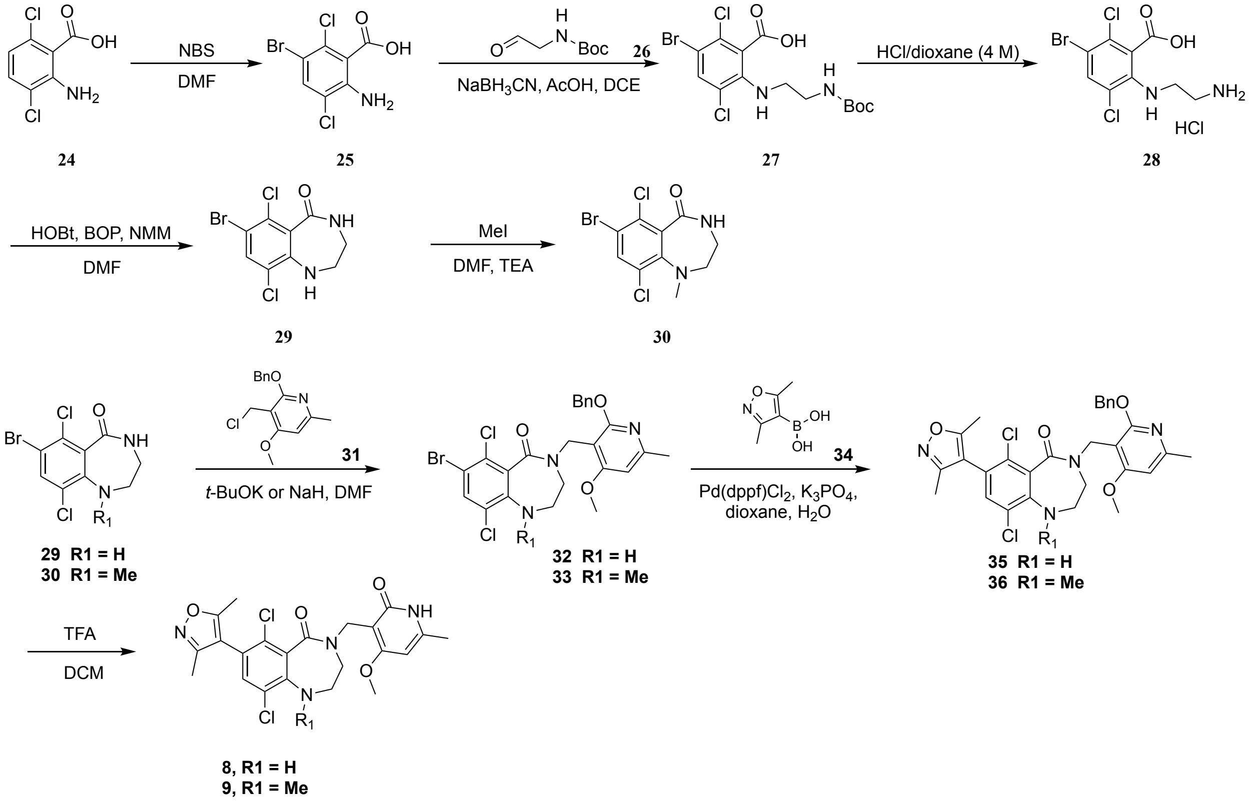


**2-Amino-5-bromo-3,6-dichlorobenzoic acid** (**25**):

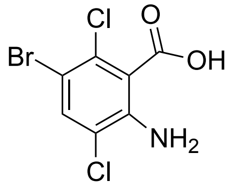

**25**

To a solution of 2-amino-3,6-dichlorobenzoic acid (**24**) (10.0 g, 48.5 mmol) in N, N-dimethylformamide (80 mL) was added NBS (9.50 g, 53.4 mmol) in portions at 0 °C. The reaction was stirred at 25 °C for 1 hour, and then added into ice-water (100 mL) causing a solid to precipitate out. The precipitate was collected by filtration. The filter cake was washed with water (2 x 100 mL) and then triturated with petroleum ether (2 x 100 mL) to afford 11.0 g (80%) of 2-amino-5-bromo-3,6-dichlorobenzoic acid (**25**) as a brown solid. $^{1}$H NMR (DMSO-$d_6$, 400 MHz) □ 7.73 (s, 1H), 5.93 - 5.76 (m, 2H).

**3-Bromo-6-((2-((tert-butoxycarbonyl)amino) ethyl)amino)-2,5-dichlorobenzoic acid *(27*)**

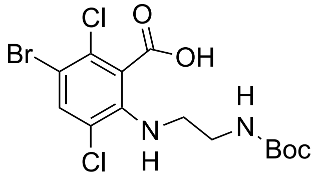


**27**

To a mixture of 2-amino-5-bromo-3,6-dichlorobenzoic acid (**25**) (5.00 g, 17.6 mmol), tert-butyl (2-oxoethyl)carbamate (**26**) (3.91 g, 24.6 mmol) and 4Å molecular sieve (5.00 g) in 1, 2-dichloroethane (50 mL) was added acetic acid (1.51 mL, 26.3 mmol) at 25 °C, and then the mixture was heated to 60 °C and stirred for 1 hour. The mixture was cooled to 25 °C, sodium borohydride (5.51 g, 87.7 mmol) was added portionwise into this. The resultant mixture was heated to 60 °C and stirred for 12 hours, then cooled to room temperature, filtered off the solid. The filtrate was added into water (50 mL) and extracted with dichloromethane (3 x 50 mL). The separated organic layers were discarded and aqueous phase was acidified by 1 M hydrochloric acid to pH~5, then extracted with ethyl acetate (4 x 100 mL), the combined organic layers were dried over anhydrous sodium sulfate, filtered and the filtrate was concentrated under reduced pressure to afford 3.30 g (44%) of 3-bromo-6-((2-((tert-butoxycarbonyl)amino) ethyl)amino)-2,5-dichlorobenzoic acid (**27**) as a yellow solid. $^{1}$H NMR (DMSO-$d_6$, 400 MHz) □ 7.59 (br. s, 1H), 7.09 (br. s, 1H), 6.54 (s, 1 H), 5.06 – 5.03 (m, 1H), 3.25 - 3.22 (m, 2H), 3.08 - 3.05 (m, 2H), 1.33 (s, 9H).

**2-((2-Aminoethyl)amino)-5-bromo-3,6-dichlorobenzoic acid (28)**

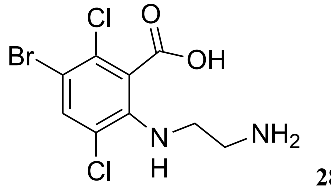


**28**

3-Bromo-6-((2-((tert-butoxycarbonyl)amino)ethyl)amino)-2,5-dichlorobenzoic acid (**27**) (3.30 g, 7.71 mmol) was added to a solution of hydrochloric acid (4 M, 10 mL) in 1,4-dioxane at 25 °C and stirred for 1 hour. The precipitate was collected by filtration, then triturated with methyl tertiary butyl ether (3 x 10 mL) to afford 3.00 g (97%) of 2-((2-aminoethyl)amino)-5-bromo-3,6-dichlorobenzoic acid (**28**) as di-hydrochloric salt as a white solid. $^1$H NMR (DMSO-$d_6$, 400 MHz) □ 8.90 (br. s, 3H), 7.85 (s, 1H), 3.43 - 3.40 (m, 2H), 2.99 - 2.95 (m, 2H). ESI (*m/z*) 328.7 [$C_9H_9BrCl_2N_2O_2$ + H]$^+$.

**7-Bromo-6,9-dichloro-3,4-dihydro-1H- benzo[e][1,4] diazepin-5(2H)-one *(29)***

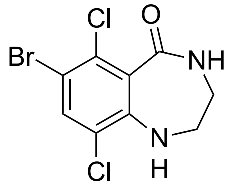


**29**

A solution of 2-((2-aminoethyl)amino)-5-bromo-3,6-dichlorobenzoic acid di-hydrochloride (**28**) (3.00 g, 7.48 mmol), BOP (4.30 g, 9.73 mmol), HOBT (1.31 g, 9.73 mmol) and 4-methylmorpholine (1.07 mL, 9.73 mmol) in N,N-dimethylformamide (250 mL) was stirred at 0 °C for 1 hour and then warmed to 25 °C and stirred for 16 hours. The mixture was poured into water (100 mL) and extracted with ethyl acetate (4 x 600 mL). The combined organic layers were washed with brine (5 x 400 mL), dried over anhydrous sodium sulfate, filtered and filtrate was concentrated under reduced pressure. The residue was triturated with ethyl acetate (2 x 5 mL), the solid was collected by filtration, and the filtrate was purified by column chromatography on silica gel (petroleum ether/ethyl acetate=1/1 to 0/1) to afford a yellow solid, which was further triturated with water (10 mL). The two solid were combined and dried in high vacuum to afford 1.20 g (52%) of 7-bromo-6,9-dichloro-3,4-dihydro-1H- benzo[e][1,4] diazepin-5(2H)-one (**29**) as an off-white solid. $^1$H NMR ($CDCl_3$, 400 MHz): □ 7.65 (s, 1H), 6.98 (br. s, 1H), 4.29 (br. s, 1H), 3.59 - 3.56 (m, 2H), 3.50 - 3.46 (m, 2H).

**7-Bromo-6, 9-dichloro-1-methyl-3,4-dihydro-1H-benzo[e][1,4]diazepin-5(2H)-one**

**(30)**

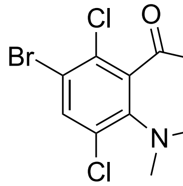

30

To a solution of 7-bromo-6,9-dichloro-3,4-dihydro-1H-benzo[e][1,4]diazepin-5(2H)-one (**29**) (0.400 g, 1.29 mmol) in N,N-dimethylformamide (4 mL) was added triethylamine (0.539 mL, 3.87 mmol), followed by methyl iodide (0.161 mL 2.58 mmol) at 25 °C. After addition, the mixture was heated to 70 °C and stirred for 16 hours. The mixture was cooled to 25 °C and added into water (10 mL). The aqueous phase was extracted with ethyl acetate (4 x 30 mL), washed with brine (3 x 20 mL), dried over anhydrous sodium sulfate, filtered and concentrated under reduced pressure. The residue was purified by column chromatography on silica gel (petroleum ether/ethyl acetate = 2/1 to 0/1) to afford 340 mg (81%) of 7-bromo-6, 9-dichloro-1-methyl-3,4-dihydro-1H-benzo[e][1,4]diazepin-5(2H)-one (**30**) as yellow oil. $^1$H NMR ($CDCl_3$, 400 MHz) □ *d* 7.68 (s, 1H), 6.31 (br. s, 1H), 3.29 - 3.25 (m, 4H), 3.04 (s, 3H). ESI (*m/z*) 324.7 $[C_{10}H_9BrCl_2N_2O + H]^+$.

**4-((2-(benzyloxy)-4-methoxy-6-methylpyridin-3-yl)methyl)-7-bromo -6,9-dichloro-3,4-dihydro-1H-benzo[e][1,4]diazepin-5(2H)-one *(32)***

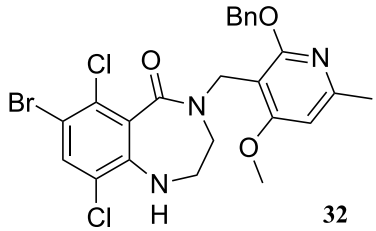

32

To a solution of 7-bromo-6,9-dichloro-3,4-dihydro-1H-benzo[e][1,4]diazepin-5(2H)-one (**29**) (0.200 g, 0.645 mmol) in N,N-dimethylformamide (1 mL) was added a solution of potassium tert-butoxide (1 M, 0.968 mL, 0.968 mol) in tetrahydrofuran at 20 °C. After stirring for 10 minutes, the mixture was cooled to 0 °C and a solution of 2-(benzyloxy)-3-(chloromethyl)-4-methoxy-6-methylpyridine (**31**) (0.179 g, 0.645 mmol) in N,N-dimethylformamide (1 mL) was added into it. Then the mixture was heated to 40 °C for 30 minutes. The mixture was added water (10 mL) and extracted with ethyl acetate (3 x 30 mL). The combined organic phase was washed with brine (2 x 20 mL), dried over anhydrous sodium sulfate, filtered and concentrated under reduced pressure. The residue was purified by column chromatography on silica gel (petroleum ether/ethyl acetate = 5/1 to 1/1) to afford 150 mg (41%) of 4-((2-(benzyloxy)-4-methoxy-6-methylpyridin-3-yl)methyl)-7-bromo -6,9-dichloro-3,4-dihydro-1H-benzo[e][1,4]diazepin-5(2H)-one (**32**) as a white solid. $^1$H NMR (DMSO-$d_6$, 400 MHz) □ 7.80 (s, 1H), 7.46 - 7.45 (m, 2H), 7.35 - 7.28 (m, 3H), 6.72 (s, 1H), 5.52 (br. s, 1H), 5.37 (s, 2H), 4.66 (s, 2H), 3.83 (s, 3H), 3.25 - 3.24 (m, 2H), 3.18 - 3.16 (m, 2H), 2.39 (s, 3H). ESI (*m/z*) 551.9 $[C_{24}H_{22}BrCl_2N_3O_3 + H]^+$.

**4-((2-(Benzyloxy)-4-methoxy-6-methylpyridin-3-yl)methyl)-7-bromo -6,9-dichloro-1-methyl-3,4-dihydro-1H-benzo[e][1,4]diazepin-5(2H)-one *(33)***

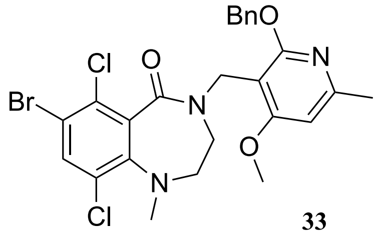


**33**

To a solution of 7-bromo-6,9-dichloro-1-methyl-3,4-dihydro-1H-benzo[e][1,4] diazepin-5(2H)-one (**30**) (0.250 g, 0.772 mmol) in N,N-dimethylformamide (2.5 mL) was added sodium hydride (60% purity in mineral oil) (0.093 g) at 25 °C. After stirring for 10 minutes, a solution of 2-(benzyloxy)-3-(chloromethyl)-4-methoxy-6-methylpyridine (**31**) (0.236 g, 0.849 mmol) in N,N-dimethylformamide (2.5 mL) was added dropwise to the mixture at 0 °C. After addition, the mixture was heated to 40 °C and stirred for 30 minutes. The mixture was added into water (60 mL) and extracted with ethyl acetate (3 x 30 mL). The combined organic phase was washed with brine (80 mL), dried over anhydrous sodium sulfate, filtered and concentrated under reduced pressure. The residue was purified by column chromatography on silica gel (petroleum ether/ethyl acetate = 100/1 to 8/1) to afford 300 mg (64%) of 4-((2-(benzyloxy)-4-methoxy-6-methylpyridin-3-yl)methyl)-7-bromo -6,9-dichloro-1-methyl-3,4-dihydro-1H-benzo[e][1,4]diazepin-5(2H)-one (**33**) as yellow oil. $^1$H NMR ($CDCl_3$, 400 MHz) □ 7.59 (s, 1H), 7.47 - 7.45 (m, 2H), 7.38 - 7.29 (m, 3H), 6.42 (s, 1H), 5.45 - 5.42 (m, 2H), 5.17 -5.14 (m, 1H), 4.58 - 4.55 (m, 1H), 3.86 (s,

3H), 3.07 - 3.04 (m, 2H), 2.97 - 2.96 (m, 2H), 2.91 (s, 3H), 2.47 (s, 3H); ESI (*m/z*) 566.0 [$C_{25}H_{24}BrCl_2N_3O_3$ + H]$^+$.

**4-((2-(Benzyloxy)-4-methoxy-6- methylpyridin-3-yl)methyl)-6,9-dichloro-7-(3,5-dimethylisoxazol-4-yl)-3,4-dihydro-1H-benzo[e][1,4]diazepin-5(2H)-one *(35)***

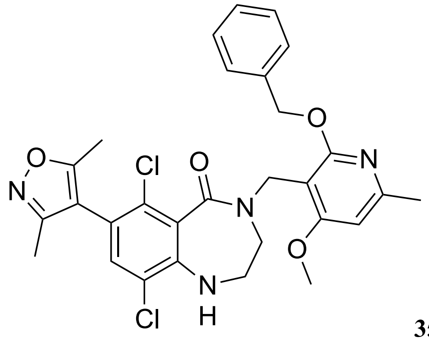


35

To a mixture of 4-((2-(benzyloxy)-4-methoxy-6-methylpyridin-3-yl)methyl)-7-bromo -6,9-dichloro-3,4-dihydro-1H-benzo[e][1,4]diazepin-5(2H)-one (**32**) (0.100 g, 0.181 mmol), (3,5-dimethylisoxazol-4-yl)boronic acid (**34**) (0.0380 g, 0.272 mmol) and potassium phosphate (0.116 g, 0.544 mmol) in 1,4-dioxane (2 mL) and water (0.4 mL) was added Pd(dppf)$Cl_2$ (0.0130 g, 0.0180 mmol) in one portion at 25 °C under nitrogen atmosphere. The reaction mixture was heated to 90 °C and stirred for 2 hours. Additional (3,5-dimethylisoxazol-4-yl)boronic acid (**34**) (0.0260 g, 0.181 mmol and potassium phosphate (0.116 g, 0.544 mmol) was added into it at 25 °C under nitrogen atmosphere. And the reaction mixture was heated to 90 °C and stirred for 3 hours. This process was repeated three times. The mixture was cooled to 25 °C and added into water (10 mL), extracted with ethyl acetate (3 x 30 mL). The combined organic phase was washed with brine (2 x 10 mL), dried over anhydrous sodium sulfate, filtered and concentrated under reduced pressure. The residue was purified by prep-TLC ($SiO_2$, petroleum ether/ethyl acetate = 2:1) to afford 50 mg (29%) of 4-((2-(benzyloxy)-4-methoxy-6- methylpyridin-3-yl)methyl)-6,9-dichloro-7-(3,5-dimethylisoxazol-4-yl)-3,4-dihydro-1H-benzo[e][1,4]diazepin-5(2H)-one (**35**) as a white solid. **1H NMR** ($CDCl_3$, 400 MHz):  7.47 - 7.45 (m, 2H), 7.36 - 7.29 (m, 3H), 7.15 (s, 1H), 6.43 (s, 1H), 5.44 (s, 2H), 4.93 - 4.86 (m, 2H), 3.86 (s, 3H), 3.32 - 3.29 (m, 4H), 2.46 (s, 3H), 2.29 (s, 3H), 2.15 (s, 3H); ESI (*m/z*) 567.1 [$C_{29}H_{28}Cl_2N_4O_4$ + H]$^+$.

**4-((2-(Benzyloxy)-4-methoxy-6 -methylpyridin-3-yl)methyl)-6,9-dichloro-7-(3,5-dimethylisoxazol-4-yl)-1-methyl-3,4-dihydro-1H-benzo[e][1,4]diazepin-5(2H)-one (36)**

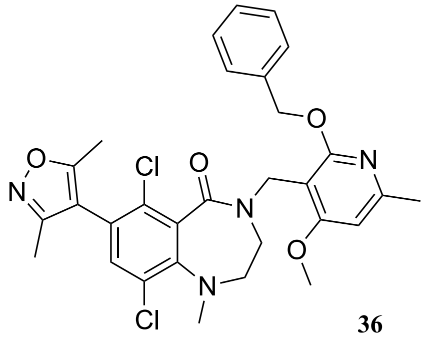


**36**

To a mixture of 4-((2-(benzyloxy)-4-methoxy-6-methylpyridin-3-yl)methyl)-7-bromo -6,9-dichloro-1-methyl-3,4-dihydro-1H-benzo[e][1,4]diazepin-5(2H)-one (**33**) (0.250 g, 0.442 mmol), (3,5-dimethylisoxazol-4-yl)boronic acid **34** (0.0690 g, 0.486 mmol) and potassium phosphate (0.282 g, 1.33 mmol) in 1,4-dioxane (1 mL) and water (0.2 mL) was added Pd(dppf)$Cl_2$ (0.0320 g, 0.0440 mmol) in one portion at 25 °C under nitrogen atmosphere. Then the mixture was heated to 90 °C and stirred for 2 hours. The mixture was cooled to 25 °C and added (3,5-dimethylisoxazol-4-yl)boronic acid (0.0310 g, 0.221 mmol) in one portion at 25 °C under nitrogen atmosphere. The mixture was heated to 90 °C and stirred for another 2 hours. The mixture was cooled to 25 °C and added water (20 mL). The mixture was extracted with ethyl acetate (3 x 40 mL). The combined organic phase was washed with brine (20 mL), dried over anhydrous sodium sulfate, filtered and concentrated under reduced pressure. The residue was purified by column chromatography on silica gel (petroleum ether/ethyl acetate = 5/1 to 1/1) to afford 150 mg (48%) of 4-((2-(benzyloxy)-4-methoxy-6 -methylpyridin-3-yl)methyl)-6,9-dichloro-7-(3,5-dimethylisoxazol-4-yl)-1-methyl-3,4-dihydro-1H-benzo[e][1,4]diazepin-5(2H)-one (**36**) as colorless oil. $^{1}$H NMR ($CDCl_3$, 400 MHz) □ 7.47 - 7.45 (m, 2H), 7.37 - 7.29 (m, 3H), 7.15 (s, 1H), 6.43 (s, 1H), 5.46 - 5.43 (m, 2H), 5.18 - 5.15 (m, 1H), 4.60 - 4.57 (m, 1H), 3.86 (s, 3H), 3.14 - 3.10 (m, 2H), 3.05 - 3.02 (m, 2H), 2.97 (s, 3H), 2.47 (s, 3H), 2.29 (s, 3H), 2.18 - 2.15 (m, 3H); ESI (*m/z*) 581.1[$C_{30}H_{30}Cl_2N_4O_4$ + H]$^+$.

**6,9-Dichloro-7-(3,5-dimethylisoxazol -4-yl)-4-((4-methoxy-6-methyl-2-oxo-1,2-dihydropyridin-3-yl)methyl)-3,4-dihydro-1H-benzo[e][1,4]diazepin-5(2H)-one (8)**

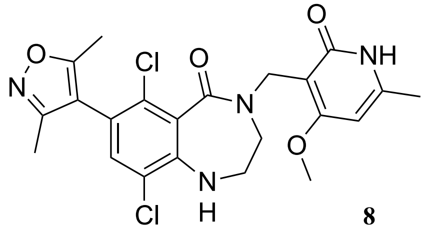

**8**

To a solution of 4-((2-(benzyloxy)-4-methoxy-6-methylpyridin-3-yl)methyl)n-6,9- dichloro-7-(3,5-dimethylisoxazol-4-yl)-3,4-dihydro-1H-benzo[e][1,4]diazepin-5(2H)-one (**35**) (0.0500 g, 0.0760 mmol) in dichloromethane (1 mL) was added 2,2,2-trifluoroacetic acid (0.834 mL, 11.3 mmol) at 25 °C. The mixture was heated to 40 °C and stirred for 2 hours. The mixture was concentrated under reduced pressure. To the residue was added 1M aqueous solution of sodium hydroxide dropwise to adjust pH~9, extracted with ethyl acetate (10 mL*3), the combined organic layers were dried over anhydrous sodium sulfate, filtered and concentrated under reduced pressure. The residue was purified by prep-HPLC (column: Waters Xbridge 150*25 mm* 5 um; mobile phase: [water (10 mM $NH_4HCO_3$)-ACN]; B%: 18% - 48%, 9 min) to afford 22 mg (57%) of 6,9-dichloro-7-(3,5-dimethylisoxazol -4-yl)-4-((4-methoxy-6-methyl-2-oxo-1,2-dihydropyridin-3-yl)methyl)-3,4-dihydro-1H-benzo[e][1,4]diazepin-5(2H)-one **8** as a white solid. $^1$H NMR ($CDCl_3$, 400 MHz) □ 11.92 (br. s, 1H), 7.16 (s, 1H), 5.96 (s, 1H), 4.86 - 4.83 (m, 2H), 4.33 (br. s, 1H), 3.89 (s, 3H), 3.55 (s, 4H), 2.36 (s, 3H), 2.29 (s, 3H), 2.16 (s, 3H); ESI (*m/z*) 477.1 $[C_{22}H_{22}Cl_2N_4O_4 + H]^+$.

**6,9-Dichloro-7-(3,5-dimethylisoxazol-4-yl)-4-((4-methoxy-6 -methyl-2-oxo-1,2-dihydropyridin-3-yl)methyl)-1-methyl-3,4-dihydro-1H-benzo[e][1,4]diazepin-5(2H)-one (9)**

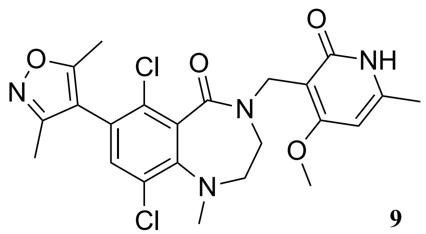

**9**

To a mixture of 4-((2-(benzyloxy)-4-methoxy-6 -methylpyridin-3-yl)methyl)-6,9 -dichloro-7-(3,5-dimethylisoxazol-4-yl)-1-methyl-3,4-dihydro-1H-benzo[e][1,4]diazepin-5(2H)-one (**36**) (0.100 g, 0.172 mmol) in dichloromethane (1 mL) was added trifluoroacetic acid (1.00 mL, 13.5 mmol) at 25°C and stirred for 16 hours. The reaction was heated to 40°C and stirred for another 1 hour. The mixture was concentrated under reduced pressure. The residue was added 1 mL methanol and an aqueous solution of sodium hydroxide (1M, 2 mL) drop wise and adjusted pH~9 causing a solid to precipitate out. The precipitate was collected by filtration. The crude product was purified by prep-HPLC (column: Waters Xbridge 150*25 mm* 5 um; mobile phase: [water (10 mM $NH_4HCO_3$)- ACN]; B%: 19%-52%, 10 min) to afford 51 mg (51%) of 6,9-dichloro-7-(3,5-dimethylisoxazol-4-yl)-4-((4-methoxy-6 -methyl-2-oxo-1,2-dihydropyridin-3-yl)methyl)-1-methyl-3,4-dihydro-1H-benzo[e][1,4]diazepin-5(2H)-one **9** as a white solid. $^1$H NMR ($CDCl_3$, 400 MHz): □ 11.83 (br. s, 1H), 7.17 (s, 1H), 5.95 (s, 1H), 5.11 - 5.08 (m, 1H), 4.55 - 4.51 (m, 1H), 3.88 (s, 3H), 3.42 - 3.41 (m, 1H), 3.28 - 3.21 (m, 3H), 3.05 (s, 1H), 2.36 (s, 3H), 2.30 (s, 3H), 2.18 - 2.16 (m, 3H); ESI (*m/z*) 491.1 $[C_{23}H_{24}Cl_2N_4O_4 + H]^+$.

### Biology

#### EZH2 Methyltransferase Assay

Compounds were screened and profiled using commercial radiometric HotSpot EZH2 methyltransferase assay service provided by Reaction Biology.

#### Cell Growth Inhibition Assay

Compounds were evaluated in cell growth inhibition assay service provided by Pharmaron.

#### H3K27me3 Flow Cytometry

293T cells were treated with either inhibitor or vehicle (DMSO) for 48hr in 90% DMEM supplemented with 10% fetal bovine serum (FBS). Cells were prepared using intracellular flow staining kit (Ebioscience). Briefly, were harvested and washed in PBS. 1mL of fixation/permeabilization working solution was added to each sample, pulse vortexed and incubated at 4°C for 60 minutes in the dark. Cells were washed twice with 2 ml of permeabilization buffer. Fluorescent conjugated H3K27me3 or Isotype control antibody (Cell Signaling Technologies) were added in 100 µl of permeabilization buffer, incubated at 4°C for 45 minutes in the dark, and then followed by two washes with 2 ml of permeabilization buffer. Cells were then resuspended in PBS supplemented with 1% FBS and acquired on an LSR-Fortessa. A ratio derived from median fluorescence intensity in cells treated at a particular inhibitor concentration relative to vehicle treated cells was calculated in order to generate dose-inhibition curves.
